# AI-Guided Resetting of Memories in Gene Regulatory Network Models: biomedical and evolutionary implications

**DOI:** 10.1101/2025.09.10.675114

**Authors:** F. Pigozzi, T. Cirrito, M. Levin

## Abstract

Molecular pathways such as gene-regulatory networks regulate numerous functions in cells and tissues that impact embryonic development, regenerative repair, aging, cancer, and many other aspects of health and disease. One important aspect of such networks is experience-dependent plasticity: their activity changes after repeated exposures to external and internal physiological stimuli. This kind of functional plasticity gives rise to habituation to pharmacological interventions (resulting in loss of efficacy over time), sensitization (resulting in unacceptable side effects after repeated use), or canalizing of undesired dynamics that recur even when the original problem has been resolved (persistent disease states). Our *in silico* analyses reveal that memories formed in gene regulatory networks can be erased by specific further experiences without any changes of network topology (leaving the connectivity in place). We present a method for discovery of stimuli that can be used to selectively delete physiological memories, which can be used to remove unwanted behaviors in biomedical and bioengineering contexts without gene therapy or genomic editing. Remarkably, not only are the training-induced gains in causal emergence not lost after stimuli that wipe memories, but we also find a positive relationship between the causal emergence and learning ability of a network, suggesting a deep asymmetry (ratchet) in the relationship between learning/forgetting and integration of collective intelligence which may have implications for evolution.

**Significance Statement:** We present an AI-driven method for discovering signals that can be used to induce physiological networks to forget specific behaviors, which can be used for applications in biomedicine and bioengineering.

## Introduction

Efforts to understand the commonalities of intelligence in unconventional embodiments – the province of the emerging field of Diverse Intelligence – benefit from model systems that are stepping stones from well-studied contexts of brains and behavior to much more exotic agents such as engineered and hybrid forms (1-7). One such model system is represented by cells and tissues; these are on the one hand mechanistically familiar (evolutionarily related) and yet alien in the sense that they exert their competencies at novel spatiotemporal scales and in difficult-to-imagine problem spaces (8). Another advantage of this model system is that progress in recognizing and learning to communicate with degrees of intelligence in living material more broadly has implications for biomedicine and bioengineering, which enables deep ideas in the Diverse Intelligence field to be cashed out as predictions and potential empirical progress (9-11). Here we focus on one such model system, the biochemical processes inside living cells.

Gene Regulatory Networks (GRNs) are a workhorse concept in biomedicine (12-14), evolutionary developmental biology (15-17), and synthetic biology (18-21). This paradigm captures the interactions of transcriptional elements – genes that regulate each other’s expression according to specific rules – but are also applicable to any kind of chemical pathway (22). Because the output of GRNs regulates biological states at the cell, tissue, organ, and whole organism level, it is important to be able to induce desired dynamics for interventions in regenerative medicine and bioengineering (23-28). These networks are known to have emergent properties (29-35), but their rational control remains challenging (36-42). We recently showed that such networks can be trained – by providing stimuli on chosen nodes and reading out responses on other nodes, well-known paradigms from behavioral and cognitive science can be used to predicably change future responses as a function of experience. Biological networks show several different kinds of learning, including habituation, sensitization, and even Pavlovian conditioning (43, 44), which are important for understanding disease states and designing strategies for intervention.

Here, we addressed the issue of pathological (undesirable) memories within GRNs and asked how they might be removed. In other words, instead of inducing new memories to be formed in molecular networks, we sought to determine whether memories might be erased (induced forgetting, as the inverse of training). Scenarios in which induced wiping of memories might be useful include habituation or sensitization to therapeutic pharmaceuticals, or unwanted associations of powerful responses to intrinsically irrelevant stimuli (45).

Pathological memories often limit the effectiveness of therapeutics, leading to increases in dosage, disease progression due to decreasing drug efficacy, severe toxicities, or the necessity to switch to more potent medications. For example, opioids are commonly prescribed and highly effective for chronic pain management (analgesia) and are particularly effective when other analgesic medications have failed, thereby serving as a last option. Morphine is a commonly prescribed opioid. Long-term tolerance to morphine can develop within days to weeks of the initiation of therapy and is often overcome by increasing the dosage (46). Dose escalation can be increased multiple-fold over the starting dose, and in some cases, up to 10-fold higher. Dose escalation also increases the risk of opioid addiction, and risks associated with morphine withdrawal. Morphine withdrawal is a life-threatening condition, often resulting from acute heart failure (47). The ability to wipe the memory of repeated administration of morphine in molecular pathway states in tissues would extend the efficacy of the starting dose, resulting in increased duration of pain management and decreased risk of addiction and death.

Rheumatoid arthritis (RA) affects 0.5-1.0% of the population and is characterized by chronic inflammation of the joints, resulting in severe pain and loss of mobility (48). There is currently no cure for RA, and patient management entails a series of increasingly more potent medications to manage inflammation. The goal of therapeutic management for RA is to maintain patients on any given drug as long as possible to delay the progression to more potent drugs which have more consequential side effect profiles. For example, corticosteroids are among the most effective drugs to manage inflammation in RA. When patients fail corticosteroids, they typically progress to biologic drugs that inhibit tumor necrosis factor (TNF). Anti-TNF drugs are highly immunosuppressive and associated with increased risk of infections and cancer. The American College of Rheumatology does not recommend corticosteroid therapy for longer than 3 months due to the side effect profile, which includes weight gain, mood changes, diabetes and immunosuppression (49). Erasing the memories of corticosteroid exposure would extend their anti-inflammatory activity, thereby delaying the administration of anti-TNF inhibitors. Moreover, wiping the memory of chronic steroid treatment would also erase the memory of the accumulated toxicities.

Proton pump inhibitors (PPIs) are commonly used to control stomach acid production to decrease the symptoms associated with gastroesophageal reflux disease or peptic ulcers and are among the most prescribed medications in the developed world (50). Long-term use of PPIs is associated with tolerance, and their discontinuation can result in hypersecretion of stomach acid, ultimately exacerbating the underlying condition. Long-term treatment also results in malabsorption, kidney failure, osteoporosis and Clostridium difficile infections (51). Wiping the memory of chronic use of PPIs would extend their efficacy and prevent toxicities, while delaying the need to switch to a different medication. With limited therapeutic options, the ability to delay a change of medication ultimately results in ensuring that patients have effective treatment options throughout their lifetime

Thus, we sought to develop a computational system to facilitate discovery of interventions that can abrogate memories in biochemical networks. We specifically examined interventions consisting only of physiological stimuli (up-or down-regulating GRN nodes via pharmacological reagents), because we seek biomedical approaches that do not require rewiring via genomic editing and gene therapy. Simulating real biological networks and training them, we examined four methods to identify memory-wiping interventions. We report a method for efficient removal of specific memories, and the remarkable finding that the causal emergence increased by training (52) is not only positively correlated with the learning ability of a network, but also not reduced the by forced memory wiping.

## Results

### A system for evaluating effects of stimuli on physiological memories

We developed a procedure to search for stimuli that reset a GRN’s memory, considering 29 real-world, peer-reviewed GRN models from the BioModels database (https://www.ebi.ac.uk/biomodels/) (53), described as continuous-time dynamical systems (i.e., a set of ordinary differential equations). While GRNs can form several different kinds of memory, *Associative* memory was the focus of our study, because it is the most complex kind of memory and models the scenario of conditioning between stimuli which has many practical applications.

Associative memory involves a triplet of nodes (a circuit): a target response R, an unconditional stimulus UCS that regulates R, and a neutral stimulus NS that does not regulate R. We first relax the GRN to allow it to settle on a steady state (*r*elaxation phase), and then we train it by stimulating UCS and NS simultaneously (training phase). If, finally, stimulation of NS only regulates R (test phase), we say the network has learned to associate NS with UCS, turning NS into a conditional stimulus CS. Out of all the circuits, 808 (among 19 networks) passed this pretest and were considered in the following. In every resetting experiment, we stimulate the network for an additional resetting phase during which we apply given stimuli and record the gene expression values at the end; if the response R is no longer up/down regulated because of the intervention, we say the memory is *reset* (see Materials and Methods).

A stimulus requires, from an intervention point of view, defining what nodes to stimulate, at what values, and when. We consequently formulated our *search cube* (depicted in Figure 2) as being inscribed by three axes:

1. The *combination* of nodes stimulated, which ranges from 1 to all the nodes.
2. Their *stimulation value*, which can be fixed or tunable.
3. The *time* at which stimulations happen, which can be only at the beginning of a simulation, or tunable.

**Figure 1.**
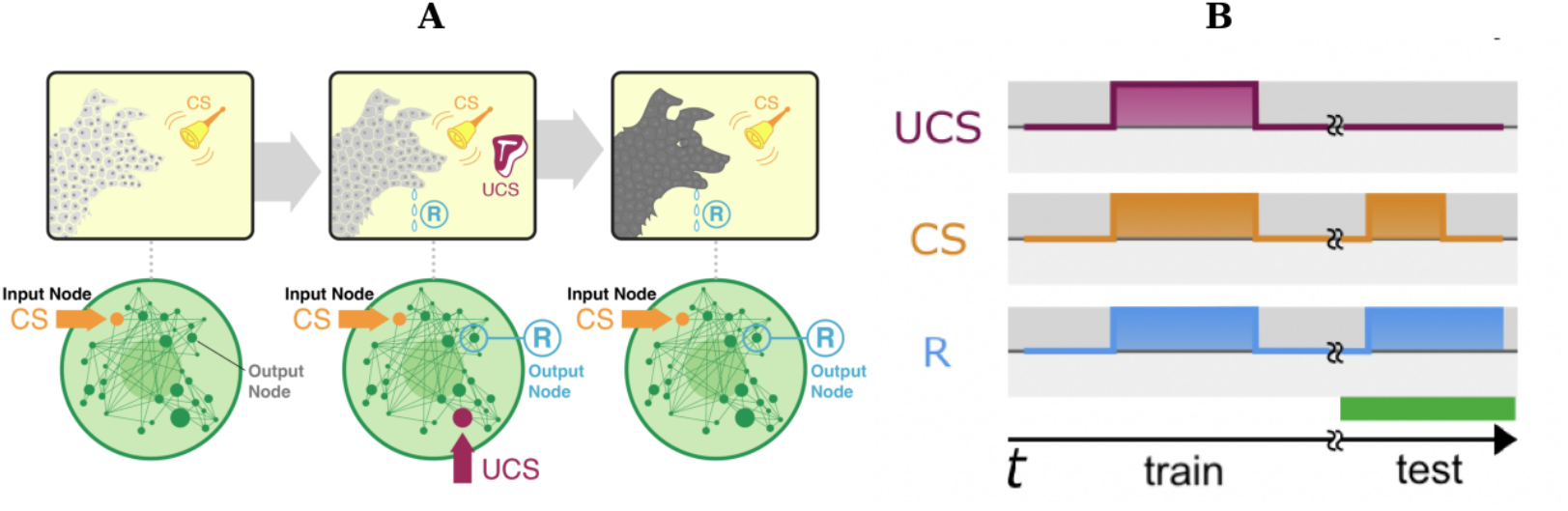
Associa8ve condi8oning in gene regulatory network models. A) Conceptual schematic of the Pavlovian conditioning training paradigm applied to gene regulatory networks. The Conditioned Stimulus (CS) is initially neutral in that it does not elicit a Response (R), while an Unconditioned Stimulus (UCS) does. For example, dogs eventually respond to the CS by forming an association between the CS and UCS after they are presented together. The same has been done in GRN pathways, by mapping the CS, UCS, and R roles to specific nodes in the network and stimulating them by temporarily raising their activity level (e.g., increasing their expression via a chemical ligand). Associative memory also raises the question of how to erase the memory, or “reset” the network to its native state, which is relevant for applications where the memory represents a clinically toxic regulation. Used with permission from [2]. B) Corresponding time-dependent levels of the genes selected from panel (A) during associative conditioning. Simulation time is demarcated on the x-axis and gene expression levels are shown on the y-axis. During the training phase, we pair a UCS and CS stimuli to regulate a response R. If associative conditioning takes place, we observe that the CS alone (i.e., with no UCS stimulation) regulates R. In the schematic, we illustrate stimulation/regulation as upticks of the expression levels over a baseline value; in reality, gene expression can have a quantitatively different shape, but the principle stays the same. Used with permission from [2].

**Figure 2:**
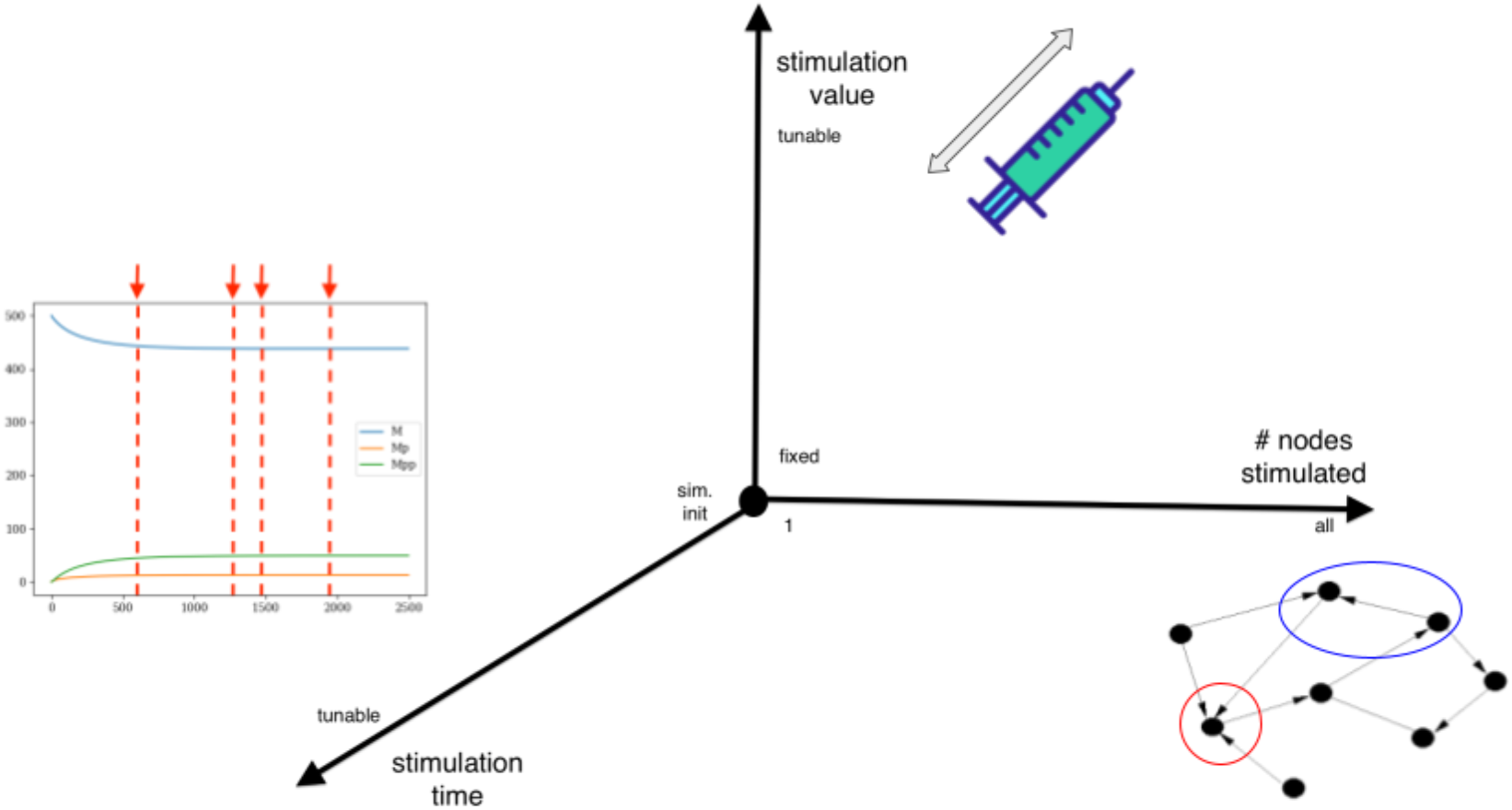
The three axes to search for interventions to control GRNs away from memories. We considered three independent parameters governing possible stimulations of a GRN, mapping them to three orthogonal axes for the space to search: the combination of nodes to stimulate, the values to which they will be set at each stimulation (corresponding to concentration of drug or transgene that activates the corresponding network node), and the time of stimulation. Since it is impossible from a computational point of view to test every possible combination of nodes, stimulation values, or times, we resort to AI to intelligently and efficiently search along the three axes.

However, it is impossible from a computational point of view to test every possible point along the three axes. For example, if we search for which nodes to stimulate (axis 1) for a network with *n* nodes, there will be 3^*(n* - 2)^ and at most *n*–2 (to exclude the R and CS) of no-stimulation/up-stimulation/down-stimulation; this number grows exponentially with the network size and becomes prohibitive even for modestly-sized networks. Hence, we resort to Artificial Intelligence (AI) to intelligently and efficiently search along the three axes.

For a given axis, the AI searches along the spectrum of values stretching from the origin to the maximum value represented by the axis. For the first axis, the combination of nodes, the AI can choose a minimum of 1 (origin) and up to a maximum of all the nodes. In the second axis, the stimulation value can be fixed (origin) or tunable; in the latter case, the AI can search for any value over the real domain of numbers. Finally, for the third axis, the time at which a stimulation happens can be only one, at the beginning (origin) up to every instant until the termination of a simulation.

### Stimulating one specific node can reset most of the memories

As a baseline, we considered the simplest treatment, corresponding to the origin in the cube of Figure 2. We stimulated only one node at a fixed, predetermined value (see Materials and Methods) and only at the first timestep of a simulation. We first searched over all the nodes by brute-force. In doing so, we excluded the R node, because it is the target we aim at stimulating by acting on the CS and UCS, and the CS node, since one can trivially break Pavlovian conditioning by ringing the bell a lot while not giving the dog any meat and the dog will eventually unlearn the association.

For 48.27% of the memories – almost half of them – our baseline finds at least one stimulus that resets that specific memory and this statistic was consistent with our previous work (54), where this treatment first appeared. Moreover, we inspected how the breakability of memories differed by network and reported the results in Figure 3. The distribution by network was uneven; in other words, different networks developed memories that were more resettable than in other networks. For example, the memories in networks #22 and #31 (the regulation of the circadian rhythm in *Drosophila melanogaster* and the MAPK cascade in *Xenopus laevis*, respectively) always reset, while those in #3 (the mitotic cell cycle of amphibians) never. We found the correlation between the fraction of reset memories per network and network to be significant with Pearson’s correlation (ρ=0.58, p<0.01). These results revealed that a slight majority of associative memories were not resettable with a brute-force approach, motivating our next experiments using AI.

**Figure 3:**
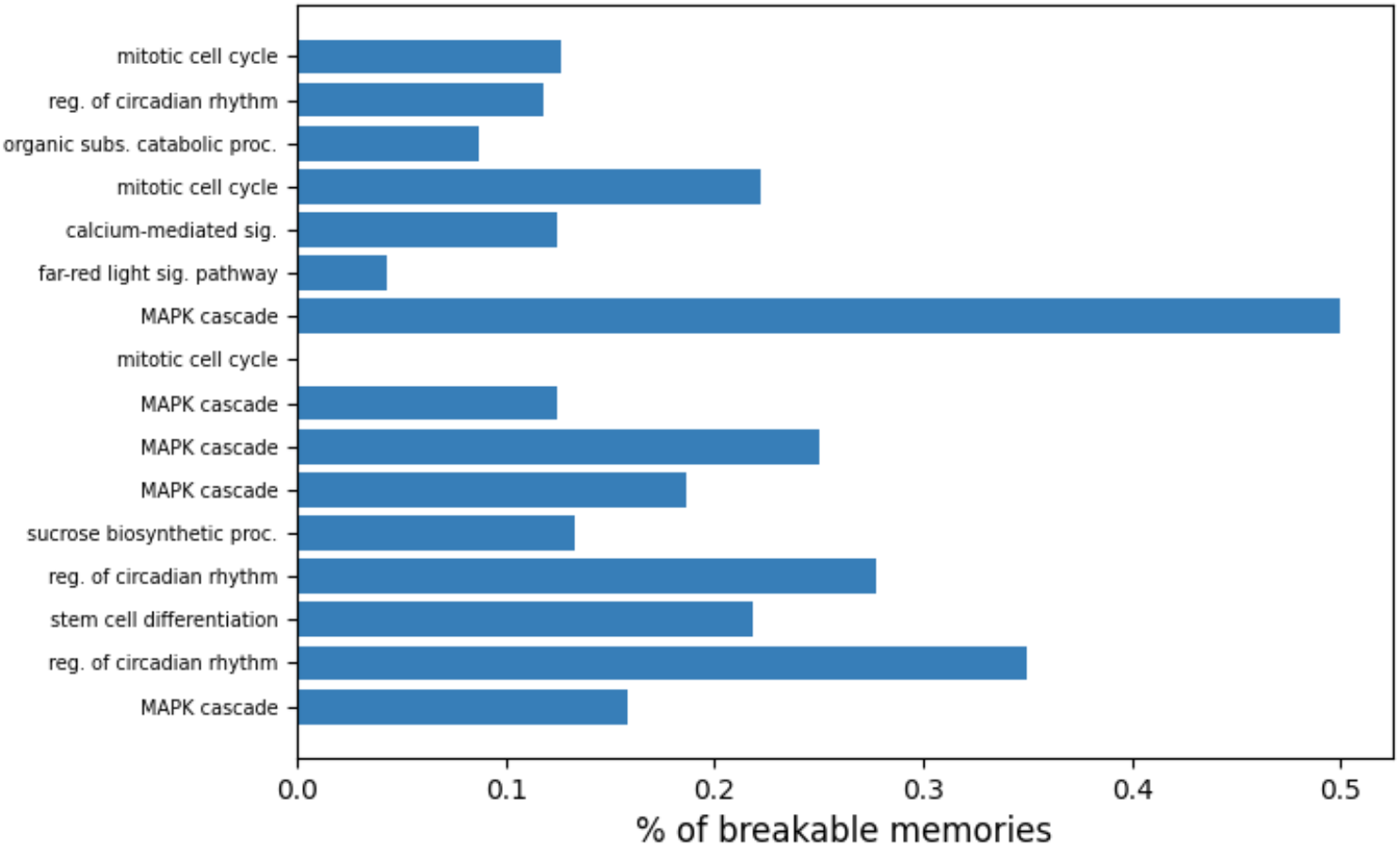
Different networks presented memories with different levels of breakability. Fraction of memories for which there existed at least one stimulation that reset them, broken down by GRN network, using brute-force search. A pattern existed: some networks presented memories that were, generally, more resettable, while others did not. The correlation between network and breakability was statistically significant.

### Optimizing along the stimulus time parameter enables erasure of the most memories

We searched over the three axes of the cube in Figure 2. The first axis (searching over the combinations of nodes to stimulate) consisted of a space whose size grew exponentially (e.g., for a network with *n* nodes, there were 3^(*n* - 2)^ possible combinations of up-stimulate/down-stimulate/no-stimulate) and was intractable for brute-force, while the second axis (searching over the nodes’ stimulation values) implied a continuous search space, not amenable to brute force since it consisted of infinitely-many points. To overcome these issues, we relied on metaheuristics (55), a family of optimization algorithms that adopt intelligent search strategies to sample a space that is too large to be explored extensively and return a solution that is, at least, a local optimum. For the first two axes, we resorted to Genetic Algorithms (GAs) (56), an established class that excels at both combinatorial (first axis) and numerical (second axis) spaces; we present a schematic in Figure 4 and a technical description in the Materials and Methods. With the third axis, the one of time, we relied on Reinforcement Learning (RL) (57) since it treats the optimization of a decision-making agent over its lifetime; we present a schematic for our RL setup in Figure 5 and provide a technical description in the Materials and Methods. We report the results – together with the previous baseline for comparison – in Figure 6, with the fraction of memories (by network) that each experiment reset, bootstrapped over 5 random seeds and after enough iterations had elapsed to guarantee convergence for all algorithms.

**Figure 4:**
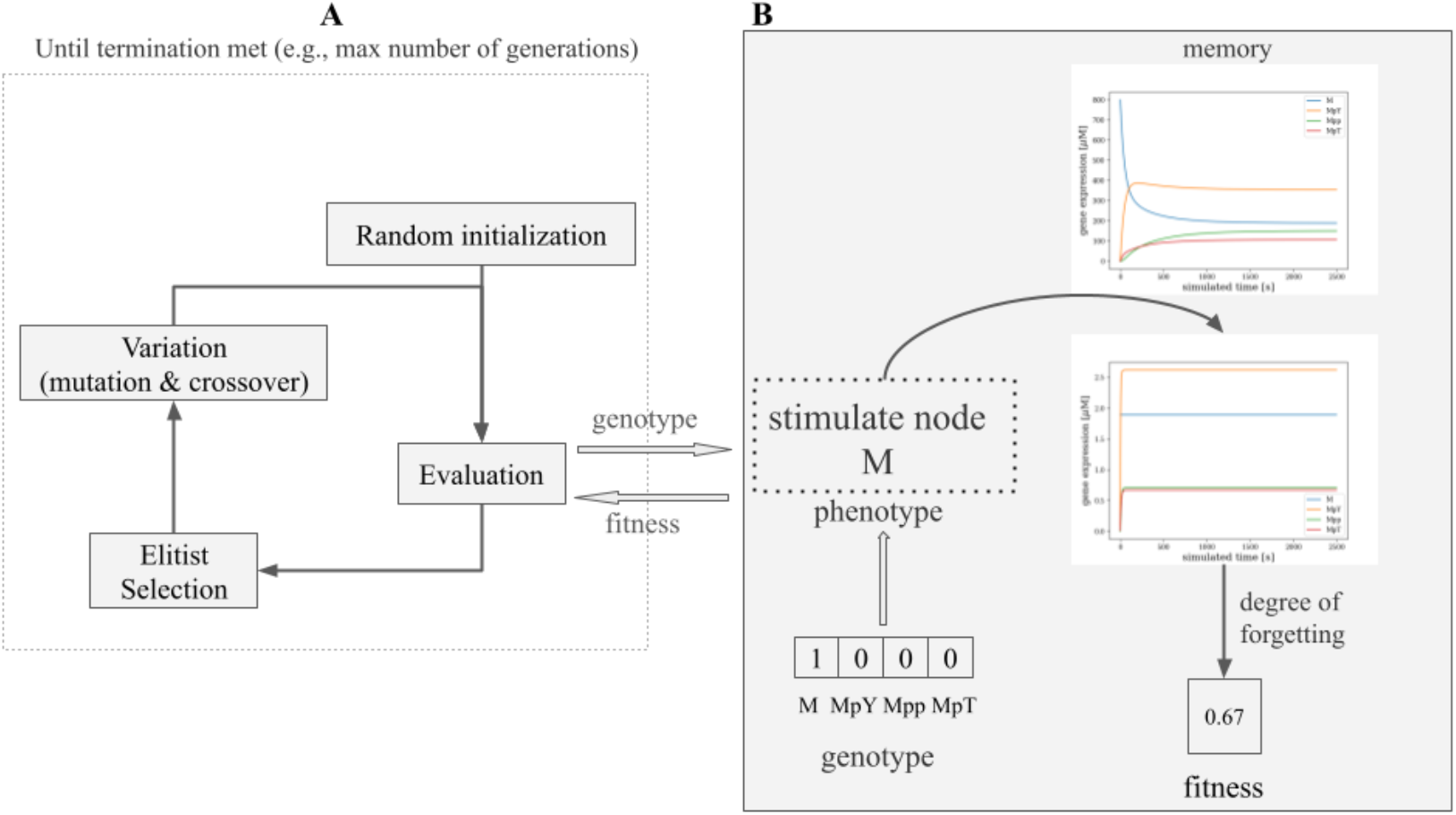
Genetic Algorithm method for discovering effective memory-blocking stimuli. A) The GA used to search the space of possible stimulation parameters. We started by randomly initializing a population of solutions; the algorithm iterated by evaluating the solutions’ fitness (see below), discarding the worst through elitist selection, varying the survivors through mutation and crossover, until a fixed number of generations had been expended. B) How evaluation took place. A solution consisted of a genotype that, for the experiment depicted here, told which nodes of a GRN to stimulate, represented as a binary string. The genotype’s translation into an actual intervention, the phenotype, followed in the sense that 1 meant stimulation and 0 meant no. We then applied the intervention and recorded the resulting gene expression levels after a simulation. The degree of memory forgetting (a real number) was then used as the fitness for the solution.

**Figure 5:**
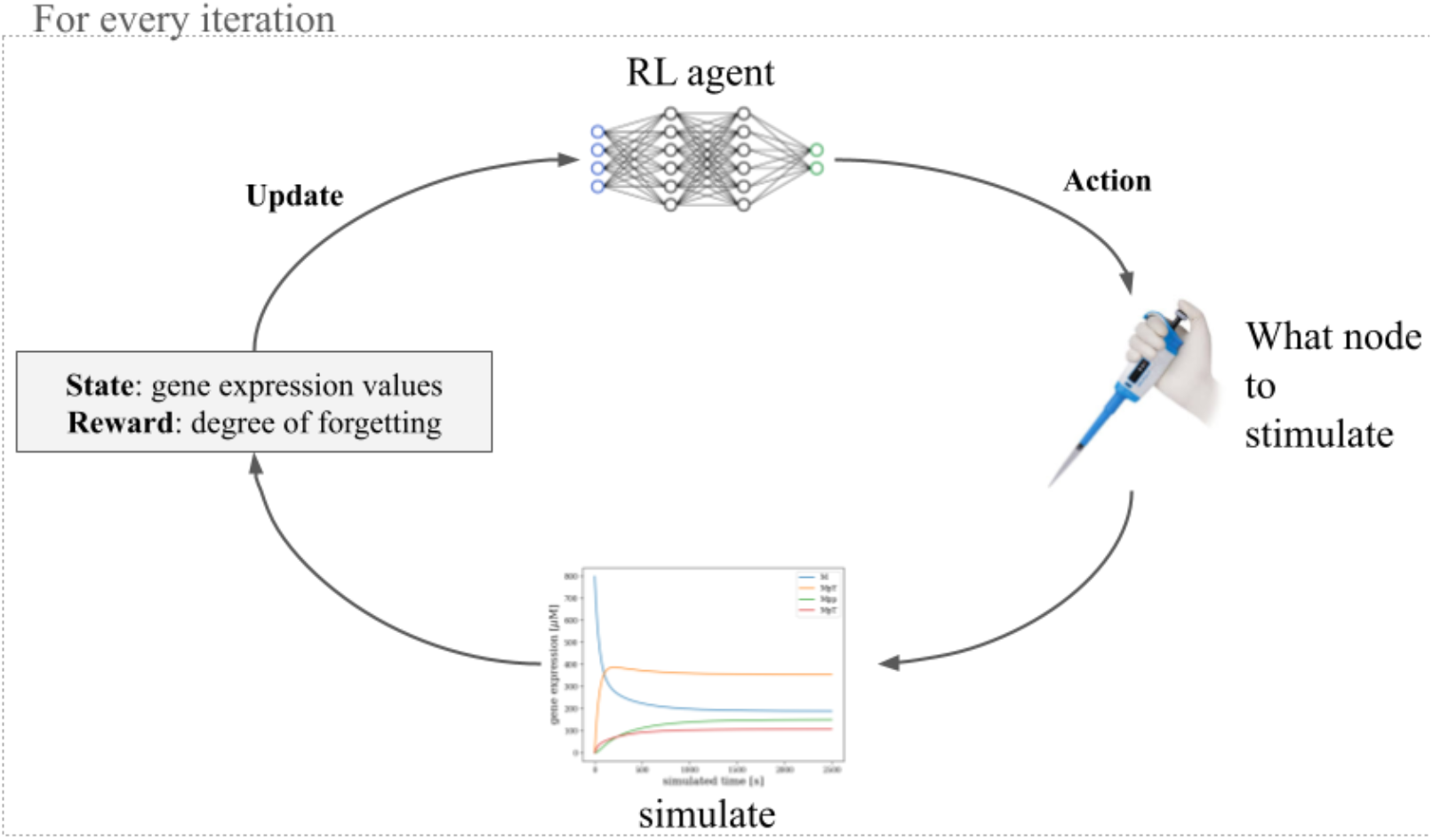
Reinforcement Learning method for discovering effective memory-blocking stimuli. An RL agent learns through trial-and-error over the course of several environment interactions. At every interaction, the agent (implemented as a neural network controller) output an action (in our case, what node to stimulate, if any), which was then applied by simulating the network for a short time and recording as the “state” its mean gene expression levels and as the “reward” the degree of memory reset. The state was then input at the next iteration, while the neural network parameters were updated to maximize the reward.

**Figure 6:**
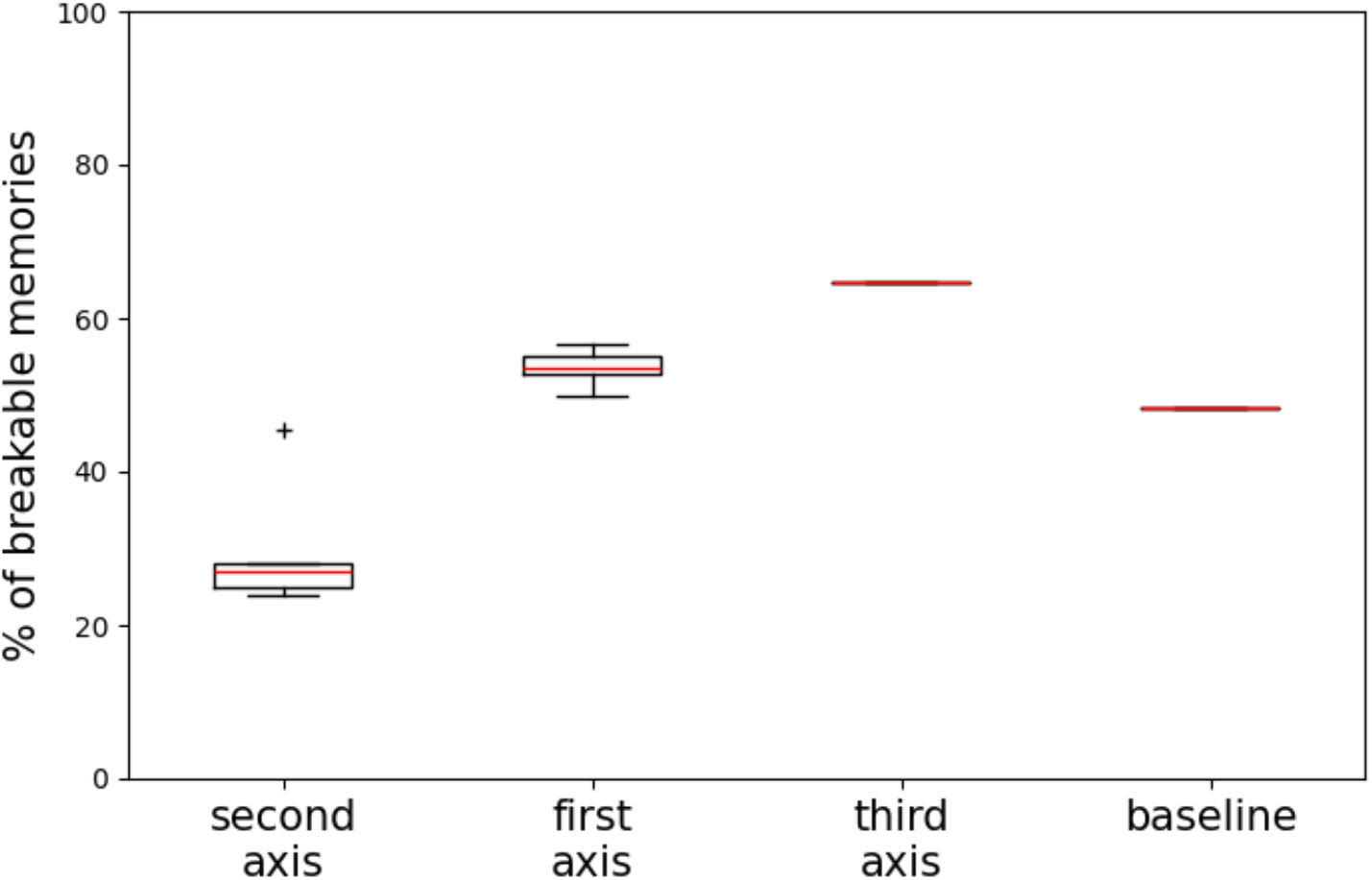
Search axes differed by the fraction of memories they reset. Fraction of memories of each network reset by each procedure; each point hence represents one network. RL – to search along the third axis – performed the best at 64.71 ± 1.18%. Results were bootstrapped over 5 random seeds. All the differences were statistically significant.

We found that RL reset the highest number of memories at 64.71±1.18%, a median improvement of 11.14% over the second best-performing approach and 16.56% over the baseline. We also found that searching over the first axis improved performance to 53.57% of the memories reset, more than half. However, the search over the stimulation values reduced performance to 26.92%. The differences among all the experiments are significant with the two-sided Mann-Whitney U test (p<0.001). These results showed that searching along the time axis is the most promising intervention for resetting memories in GRNs.

### Breakability of the memories depends on the biology and the properties of the network

We analyzed the relationship between the breakability of memories and the biology of the networks, in particular, the taxon and gene ontology of the biological context for each network. After retrieving this information from the database project source, we visualized the relationship in Figure 7 with the fraction of reset memories, according to the different categories. Breakability differed by taxon and gene ontology: some taxa (e.g., plants) and ontologies (e.g., MAPK cascade) yielded more resettable memories, while others (e.g., regulation of the circadian rhythm) did not. We found no obvious relationship between the susceptibility to memory erasure and the evolutionary position of the network, but all of them were within the range of 38.96% and 68.44%.

**Figure 7:**
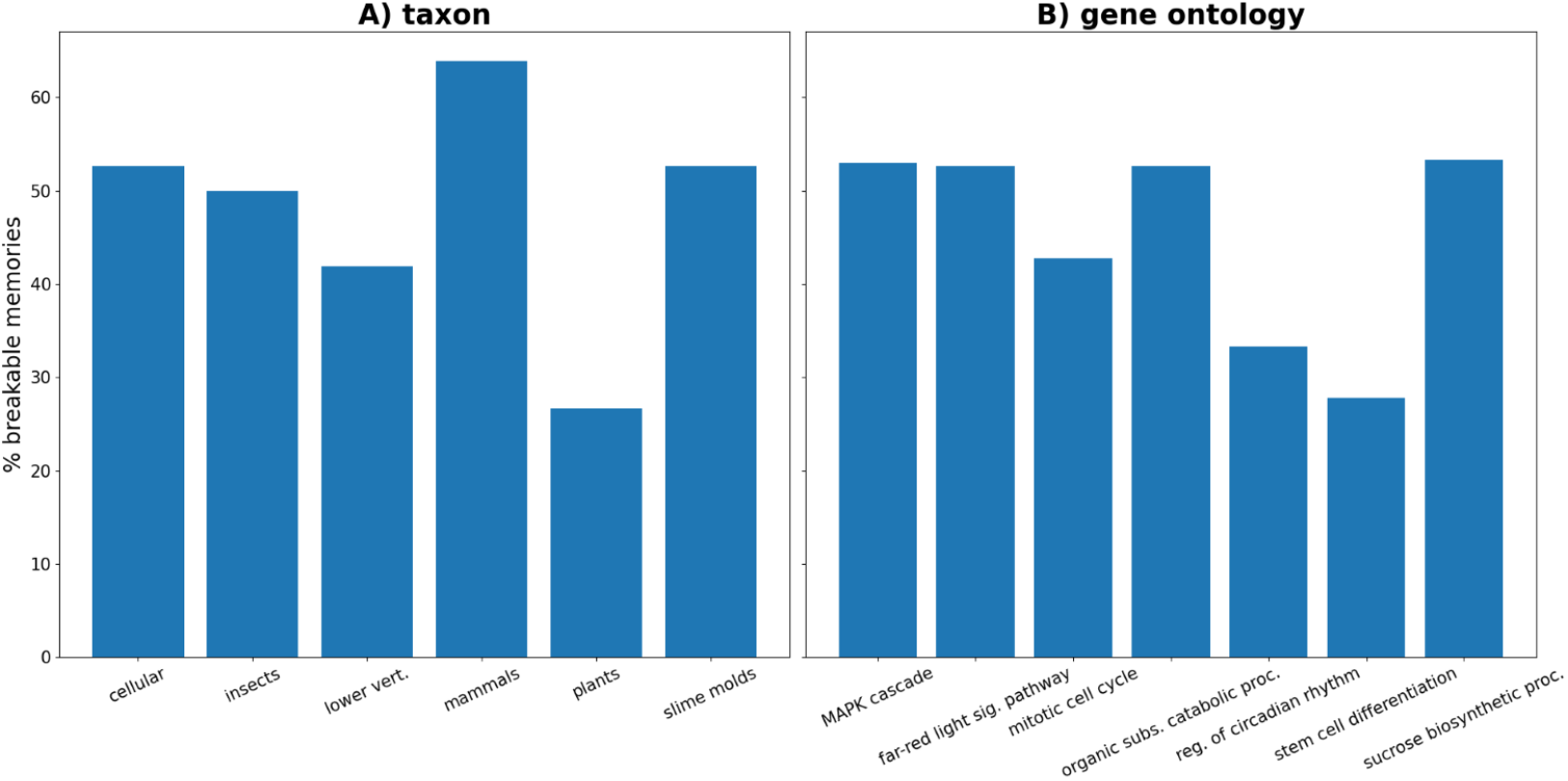
Different taxa and gene ontologies differed by the breakability of memories. Fraction of reset memories according to the biology of the network, in particular the taxon (A) and the gene ontology (B). Breakability differed by taxon and gene ontology: some taxa (e.g., plants) and ontologies (e.g., MAPK cascade) yielded more resettable memories, while others (e.g., regulation of the circadian rhythm) did not.

We next looked for relationships between the memory properties of networks and their topology metrics. We tested the correlation between the number of reset memories of a network and several network (number of nodes, edges, in-degree, out-degree, betweenness centrality, PageRank, and HITS) and dynamical systems (correlation dimension, Lyapunov coefficients, Hurst coefficient, Detrended Fluctuation Analysis: DFA) theoretic properties. We found a significant positive correlation between breakability and the number of edges (with the Pearson’s correlation test, ρ=0.55, p=0.0257); peculiarly, the same there was no significance with the number of nodes, pointing out that what mattered was the network path density, not the network size. We conjectured the reason to be more possible pathways to exert control along. Finally, we remark that we found an almost significant negative correlation with the DFA score (with the Pearson’s correlation test, *ρ* =-0.47, p=0.06), which measures the self-affinity of a trajectory (58); in other words, the higher the DFA score, the more long-range correlations in the gene expression profiles over time, the less resettable the memories of a network were. The reason may have been that more long-range correlations make a GRN more robust to external perturbations like interventions.

### Resetting memories increases the causal emergence of the network

Previous research has shown that training GRNs for associative memory increases the causal emergence of the network (52): how much information the “whole” (the network, as an integrated agent) provides about the future evolution of the system that cannot be predicted by the single parts (the nodes) only. Following (52), we measured causal emergence using the Φ *ID*decomposition, an established methodology for temporal data (like our gene expression trajectories). Causal emergence is a numerical quantity (measured in natural units) such that the higher, the more emergent (or integrated) a system is; we refer the reader to the original literature for the full mathematical derivation (59, 60).

For simplicity, we considered the baseline treatment (origin of the cube of Figure 2) and computed the causal emergence during the test and reset phases for each stimulus of the baseline that successfully reset the memory. We found that causal emergence increased from the test to the reset phase in 63.95% of the cases, or 249 out of 390. Having verified the causal emergence increases significant with the one-sided Wilcoxon test for paired samples (p<0.001, with Bonferroni correction), we concluded that resetting a memory, on average, increased the causal emergence of the agent. Crucially, for the stimuli that did not reset the memory, the change in causal emergence from test to reset phase was not significant (Wilcoxon test). The distributions for these two cases (stimuli that reset and stimuli that don’t) are shown in Figure 8. Thus, not only do associative memories reify an agent’s integration, but this integration is retained by the GRN and even potentiated after wiping out of the memory in a majority of cases.

**Figure 8:**
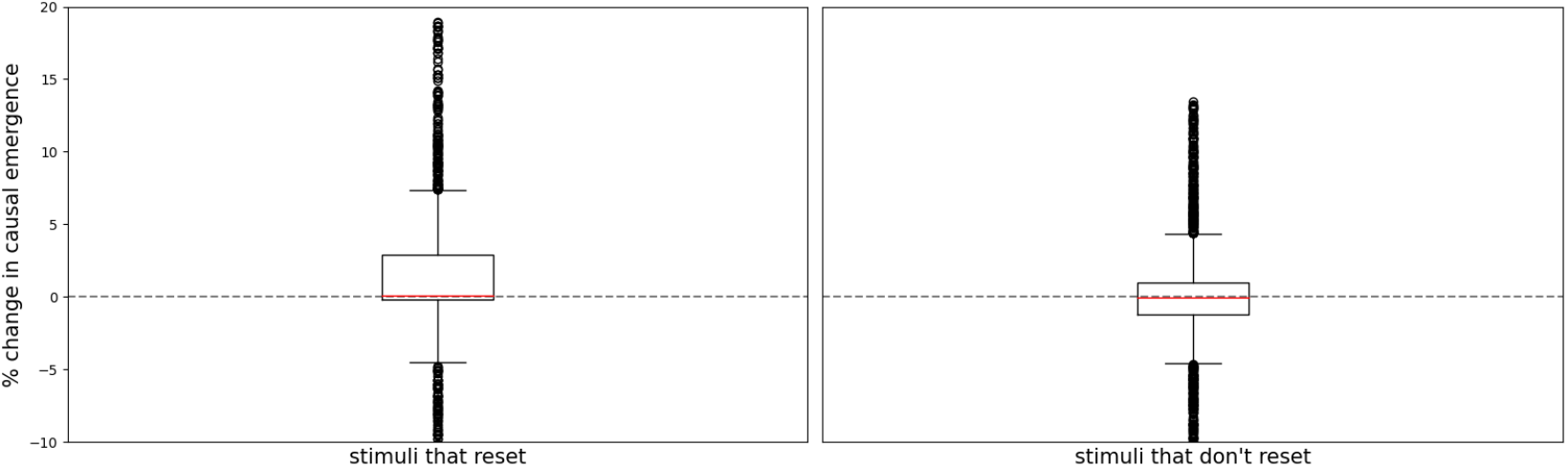
Percent change in causal emergence from the test to the reset phase for the stimuli of the baseline treatment that do reset (left) and do not reset (right) the memory. Stimuli that reset the memory have, on average, a higher increase in causal emergence.

The asymmetry in rise of integration between learning and forgetting suggested a possible feedback loop, which would be closed if higher levels of integration were themselves facilitative of learning. We found a strong relationship between the causal emergence of a network and its ability to learn. Indeed, there was a statistically significant positive correlation (p<0.001, with Pearson’s test) between the causal emergence of a network and the % of all triplets of nodes that passed the associative conditioning test (see Materials and Methods), meaning that the more causally emergent networks were also the most responsive to associative conditioning. In particular, *ρ* =0.76, indicating (together with the p-value) a strong positive correlation and, by fitting a linear regression model, we found that, on average, a 1 nat increase in causal emergence corresponded to a 35% increase in the number of a network’s triplets that passed the associative conditioning test. Similarly, a 1% increase in the number of memories led to, on average, a 1.6 nat increase in causal emergence. But, when considering the resetting instead of the learning of memories, there was no significant correlation (p>0.05 and *ρ* =0.15, with Pearson’s test) between the causal emergence of a network and the % of all memories that were reset. Thus, higher-causal emergence networks were better at learning, but not at forgetting.

## Discussion

Training GRNs for associative memory as a kind of drug conditioning has been so far possible in silico and is a promising method with which to enhance our ability to use drugs in biomedical and bioengineering contexts. The goal is to associate the effects of an inert drug with those of a powerful one, eliciting the desired response in scenarios where the powerful reagent must be limited in its use. Yet, how to *dis*associate a conditioned drug from its response (in other words, reset the memory) remained unclear. We showed that resetting these memories is indeed feasible: a simple brute-force approach reset almost half of the memories investigated, while the best of an array of AI approaches (based on GAs and RL) reset 64.71±1.18%.

Among the AI algorithms envisioned, RL, which learns how to optimally control a network over time, delivered the best results; ideally, the RL agent learned to exploit the dynamic nature of the GRN to achieve its reset goal, proving the importance of experiences for top-down control of biological networks and the relevance of adopting tools from the behavioral sciences (11). Conversely, a GA evolving a list of stimulation values performed the worst; we suspected the reason to be that enlarging the search to the stimulation values entails a more rugged fitness landscape for the evolutionary optimization. Finally, the fact that memory circuits associated with the MAPK cascade displayed higher breakability is relevant, since its central role as a memory binding subunit (like cells and molecular networks) (61-63) toward common purpose in multicellular organisms (64) may have made the network dynamics more malleable and fluid. Contrarily, the regulation of the circadian rhythm yielded the least resettable memories, as if its cyclical nature made the network dynamics less susceptible to disruptions like interventions.

There are limitations with the current work. The first and foremost concerns the need to test this method in real cells (65). Using machine learning to closed-loop control bioelectronic actuation in living cells (66), predict the response to an optogenetic system in *Escherichia coli*, and adaptive patient interventions (67), are already possible (68). Indeed, our setup provides a convenient sandbox to control models of real GRNs, but it is possible that spurious behaviors may arise in vivo. The main reason for this is that our simulated GRNs live in isolation and do not interact with the noisy and rich cell environment they are a part of, which includes, among others, bioelectric and biomechanical networks. Ongoing work will examine how, for example, the extracellular matrix’ sending and receiving signals with the GRN impacts the results. Our model does not include the ability of the GRN to also store state memory in surrounding microenvironment (other GRNs and subcellular components such as the cytoskeleton); the potential for such stigmergic (69-71) effects across molecular subsystems will be explored in future work. Moreover, not all portions of the search cube may be feasible to implement from a practical point of view. Our study provides an in silico springboard for controlling GRNs and other kinds of pathways and paves the way for future in vivo experiments, including ways to target the minority of memories that we could not erase, and similar approaches applied to other (non-associative) kinds of memories.

In addition to testing these ideas in vivo, we will scale up the capabilities of our AI algorithms. Leveraging the recent successes of foundation models (72-74) – large neural networks trained on diverse and extensive datasets – across several domains of the life sciences (75, 76), we speculate that a foundation model trained on the vastity of dynamic data on GRNs (across all strata of life) (77) will achieve generalization to a plethora of downstream tasks including, but not limited to, universal control of GRNs towards desired phenotypic states. In general, we suggest that computational (and especially AI-driven) tools to identify physiological stimuli that have desired effects is a rich area for future development, paralleling (and eventually perhaps exceeding) the plethora of efforts in drug discovery and genomic editing (9, 11).

Finally, it should be mentioned that the approach described here has broader implications beyond biomedicine. Note that it differs significantly from conventional approaches to this problem which uses methods from dynamical systems theory and similar tools to establish direct, specific molecular states by targeting specific nodes and editing the network topology. In other words, a re-wiring approach that seeks to micromanage relevant states of the complex system. We have previously argued that complementing this approach is an emerging area of life sciences – diverse intelligence research (78-86) - which leverages tools of behavioral science. By borrowing powerful paradigms from neuroscience (such as those involved in associative conditioning (87)) and applying them to domains that are not brains, new kinds of control may become possible (10, 36, 88). In this approach, the bioengineer or regenerative medicine worker try to hack the system at the level of *stimuli*, not micromanaging or forcing functions, which will have desired events as they do in a cognitive setting. For the same reason that humans could control dogs and horses thousands of years before knowing anything about brain mechanisms, we may be able to achieve very complex system-level outcomes in scenarios where direct micromanagement is not possible or not desirable.

There are a few other aspects at the intersection of this work with cognitive science that deserve mention. First, it should be noted that the space of such stimuli that is being searched by our AI system is isomorphic to the space of *hacking signals* that evolution searches. One way to see evolution applied to the multiscale competency architecture of living material (89) is as not only a search for structural solutions and hardwired mechanisms in a phenotype space, but also as a search for subtle signals (inducing training, memory-wiping, reward, etc.) that subsystems can efficiently use to control the behavior of other such subsystems and their own parts. Evolution gains as much from exploiting the competency of subsystems via simple, low-effort stimuli (not rewiring) as do scientists and physicians. Thus, a better understanding of what causes molecular pathways to change behavior (learning and forgetting) is likely to shed light on the evolution of both GRNs and other signaling mechanisms that can stimulate them.

Second, the associative nature of the training demonstrated in GRNs is, in effect, a molecular placebo – stimuli that change the way the network *interprets* future stimuli. It remains to be seen how much of the fascinating work on placebo effects (90-94) ends up facilitating physiological research at the cell level, or conversely, how the understanding of plasticity at the molecular level leverages our understanding of such plasticity at the neuro-behavioral level. Other avenues for future work include the search for disorders of memory and placebo capacity in molecular networks and an examination of whether they might underlie any known disease state.

Finally, our examination of causal emergence during learning (52) and during forgetting (above) shows an interesting phenomenon: in many networks, training forms two memories – a conventional one and a sort of meta-memory. Conventional memory is the change in behavior that was induced – for example an association. This memory disappears after induced forgetting. But the fact that the rise in Φ|ID does *not* disappear after forgetting reveals that stimuli induce a kind of memory that persists after forgetting wipes away the behavioral effects of specific learning. It is a kind of memory that is even more subtle than the distributed dynamical system changes that underlie learning in hardware-locked networks - a stable change of causal emergence metrics. Future work will determine whether this has anything to do with the known phenomenon of implicit memories in cases of anterograde amnesia (95-97). Numerous interesting connections exist between phenomena of memory in networks and aspects of cognitive science, and additional tools of that field (98-100) should be brought to bear on molecular and physiological networks to identify novel frontiers.

Finally, in the context of diverse intelligence and basal cognition, these data suggest an intriguing hypothesis relevant to the search for an explanation for the apparent generic increase in intelligence across evolutionary time and the initial origin of both, learning capacity and causal emergence of higher-level virtual governors in minimal media. Our data show that for some, even relatively small, networks, learning increases integration while integration facilitates learning. Together with the fact that forgetting does not drop the integration gains of prior experience, this forms a positive feedback look pointing to a general increase of learning capacity (a basic component of cognition). This suggests a possible mechanism for kickstarting an “intelligence ratchet” as previously observed in evolution of multicellular forms (89): if applicable to pre-biotic chemical reactions, learning and causal emergence may leverage each other even before biological evolution begins in earnest. In other words, this amazing property is not dependent on any facts of physics nor on selection but rather is a linked set of properties of mathematical objects. As such, it may be a really interesting and novel example of how biology exploits not only physics and evolutionary history but universal mathematical patterns.

Moreover, it should be noted that the property we found is not specific to gene-regulatory networks or even to biochemical pathways but could be applicable to any system whose output can be numerically measured or observed and in which interventions are possible. Relevant examples could include the economy, weather, social networks, and astronomical/cosmological structures at various scales. As with our approach to GRNs, the memory-related phenomena do not require modification of the weights of the network, which further broadens the potential relevance. Future work will examine biological and other networks at different spatiotemporal scales for the potential to support not only dynamical system learning but also learning-integration ratchets. Thus, tools for testing, and datasets comprising, unexpected instances of learning and emergent causation at higher levels of organization may be a useful component of discovery in the field of Diverse Intelligence and basal cognition (82, 83, 101, 102). Interaction with diverse intelligences in the space of possible embodied minds – those with whom we share evolutionary lineage and the impending beings beyond it – depends on the development of tools and conceptual frameworks that help recognize aspects of cognitive functions in unfamiliar guises.

## Materials and Methods

### Biological models and simulation

We curated a dataset of 29 peer-reviewed GRNs from the BioModels open-source database (53) encompassing all the strata of life (from bacteria to humans), the same adopted in (54). Each model describes a GRN whose nodes may be proteins, metabolites, or genes and whose edges are mutual relations like reactions. Networks are modeled over time according to Ordinary Differential Equations (ODEs), and so each one is a continuous-time dynamical system. Nodes take values on a continuous domain (like gene expression levels). Simulation took place in AutoDiscJAX (https://github.com/flowersteam/autodiscjax), which is not only written in the highly efficient and compute-optimized JAX language but also enables interventions on GRNs like applying stimuli, after parsing System Biology Markup Language specification files into JAX programs with SBMLtoODEjax (103). For all experiments, simulations were integrated with the fourth-order Runge-Kutta method using a step size of 0.01.

### Memory evaluation

Of the types of memory identified in (54), here we focused on associative memory, since it associates effective but toxic drugs with “placebo” triggers (104). Associative memory is analogous to Pavlovian conditioning in animals. We present a schematic in Figure 1. Associative memory involves a triplet of nodes (a *circuit*): a target response R, an unconditioned stimulus UCS that regulates R, and a neutral stimulus NS that does not. We first “relaxed” the GRN to allow it to settle on a steady state and have a baseline for its pre-training behavior (*relaxation* phase) by simulating it for *t*_s_ timesteps. We then trained it by stimulating UCS and NS simultaneously (*training* phase) for another *t*_s_ timesteps. If, finally, stimulation of NS only regulated (*testing* phase) for other *t*_s_ timesteps, we said the network had “learned” to associate UCS with NS, turning NS into a conditioned stimulus CS. Finally, during the resetting experiments, we applied the candidate stimuli (to reset a memory) during the simulation of an ulterior *resetting* phase of *t*_s_ timesteps. After preliminary experiments, we found *t*_s_=250,000 (2500 seconds of simulated time) to be sufficient for all the networks to settle on steady values.

For each network, we tested every possible triplet of nodes for associative memory. Since biological entities take on continuous values in ODE networks, we can either up-stimulate (increase the value to some extent) or down-stimulate (decrease the value up to some extent) for a specific node. Similarly, stimulation can up-regulate or down-regulate R, depending on whether stimulation increases or decreases its value. Following (54), we up-stimulated and down-stimulated by setting the quantity of a stimulus ST to *e*_max_^ST^ × 100 and *e*_min_^ST^/100 respectively, where *e*_max_^ST^ and *e*_min_^ST^ were the maximum and the minimum values the nodes attained; whenever stimulation happens at a “fixed” value, we mean these. We called R up-regulated if the mean value during testing was at least twice that during relaxation, and down-regulated if it was no more than half. Consequently, we called a memory “reset” if the value at the end of the resetting phase was less than twice the mean value during relaxation in the case of up-regulation, and more than twice in the case of down-regulation. Under this light, we define the *degree of forgetting* of an intervention on a memory as the ratio between the mean value of R during testing and the value of R at the end of the resetting phase (the higher, the more reset the memory is) in the case of up-regulation; the reciprocal in the case of down-regulation. In line with (54), we found these values for stimulation and regulation to result in associative learning in our network and simulate the delivery of real-world drugs.

Stimulation through the delivery of real-world drugs does not take place in one persistent bout, but in several time-delayed dispensations. For this reason, we applied stimulation in pulses: we partitioned the phase (whether training, testing, or resetting) into five equally sized time intervals and alternated between applying the stimulus (i.e., at the first, third, and fifth intervals) and not applying it. When a stimulus was not applied, the nodes followed their dynamics as dictated by the network. Of all the triplets of the 29 biological networks considered, the 808 that passed the pretest belonged to 19 networks (more than half), in line with (54). We considered these circuits for all the analyses.

### Genetic algorithm

We adopted a (*μ*+ *λ*) selecto-recombinative GA (56) as metaheuristic for the first two axes of the search cube (see Figure 2), in two different implementations: combinatorial (first axis) and numerical (second axis). Despite its vanilla nature, this variant has proven competitive in challenging tasks like, for example, training of virtual agents (). We depicted a general schematic of our GA in Figure 4 with the components that are common to both implementations. At the first generation, a population of *n*_pop_ individuals was randomly initialized and the fitness values, corresponding to the degree of forgetting over the simulation of a GRN resetting phase, were recorded. Then, the worst half of individuals was discarded (elitist selection) and, until *n*_pop_ children were born, a child was created with probability *p*_cx_ through crossover and probability 1 – *p*_cx_ through mutation, with its parents (two for crossover, one for mutation) selected from the survivors with tournament selection of size *n*_tour_. This procedure repeated until *n*_gen_ generations had elapsed. In the following, we described where and how the two implementations differed.

In our combinatorial GA, the genotype consisted of a string of *n* genes – one for each node – for a network of *n* nodes. Each gene took values in {0, 1, 2} and described if and how to stimulate the corresponding node: 2 for no stimulation, 0 for up-stimulation, and 1 for down-stimulation. Initialization sampled uniformly from the same integer set. Mutation flipped a child gene into one of the other integers (uniformly chosen), while crossover was one-point, meaning that each child gene had a ½ probability of inheriting from the first parent and a ½ probability of inheriting from the second parent.

In our numerical GA, the genotype consisted of a string of 2*n* genes – two for each node – for a network of *n* nodes. Each gene took values in the range [0,6] (since gene expression values cannot be negative and we found higher ranges to be unrealistic). When stimulating the GRN, for the i-the node having gene values *g*_*1*_ and *g*_*2*,_ we up-stimulated if *g*_*1*_ > 0 (down-stimulated otherwise) at the value *e*_max_^g^ x exp(*g*_*2*_) (or *e*_min_^g^ x exp(*g*_*2*_) when down-stimulating), where we took the exponential because some genes had exponentially larger ranges than others (105). Mutation perturbed the value of a uniformly chosen child gene with Gaussian noise of mean 0 and standard deviation 0.1 (clipped to [0,6]), while crossover operated in the same manner as the combinatorial GA. After preliminary experiments and relying on our expert knowledge, we set *n*_pop_=100, *p*_cx_=0.8, *n*_tour_=5, and *n*_gen_=100 for both implementations.

### Reinforcement learning

We adopted an RL approach for the third axis of the search cube (see Figure 2). Indeed, RL is the go-to method for decision problems (in our case, what interventions to apply) that span the lifetime of a dynamical system (a simulated GRN here) (57). We presented a high-level description of the RL loop in Figure 5. The agent was a “virtual” GRN controller (implemented as a neural network) that iteratively read a state from an environment (the GRN itself, represented by the gene expression levels at a given time step) and, at each iteration, chose the action that maximized the cumulative reward (the degree of memory forgetting) over time. Neural network parameters were updated over time to adjust the agent behavior towards the best actions given the state. Possible actions were which node to stimulate (at a fixed value), if any, and the reward is how “reset” is the target memory.

Given that our model of the environment consisted of non-trivial ODEs that cannot be solved analytically but must be numerically simulated, we resorted to model-free RL, which (contrary to model-based) does not assume a tractable mathematical model of the environment (57). Among all the possible choices, we selected Proximal Policy Optimization (106), an established model-free RL algorithm, which has delivered state-of-the-art results in, for example, game-playing.

We used the python implementation of r-ppo found in stable-baselines3 (107), with parameters and neural architectures (a recurrent neural network, to exploit time dependencies) set to their default values. The agent neural network took as input the average gene expression levels for all the nodes at the previous timestep and applied the softmax function (whose largest output indicated the node to stimulate at that timestep, plus one output for no node) to the output layer. A requirement for this class of methods, we normalized the reward by subtracting the running median and dividing it by the running standard deviation. Finally, we set the duration of each environment interaction to 100 timesteps (1s of simulated time), since it provided a reasonable trade-off between computational requirements and precision in the gene expression levels.

## Acknowledgements

We thank Mayalen Etcheverry, Patrick Erickson, and Santosh Manicka for help with the software and useful discussions. M.L. gratefully acknowledges support via grant 62212 from the John Templeton Foundation, via grant TWCF0606 from the Templeton World Charity Foundation, and via a sponsored research agreement from Astonishing Labs.

